# Manifold alignment reveals correspondence between single cell transcriptome and epigenome dynamics

**DOI:** 10.1101/130336

**Authors:** Joshua D. Welch, Alexander J. Hartemink, Jan F. Prins

## Abstract

Single cell genomic techniques promise to yield key insights into the dynamic interplay between gene expression and epigenetic modification. However, the experimental difficulty of performing multiple measurements on the same cell currently limits efforts to combine multiple genomic data sets into a united picture of single cell variation. We show that it is possible to construct cell trajectories, reflecting the changes that occur in a sequential biological process, from single cell ATAC-seq, bisulfite sequencing, and ChIP-seq data. In addition, we present an approach called MATCHER that computationally circumvents the experimental difficulties inherent in performing multiple genomic measurements on a single cell by inferring correspondence between single cell transcriptomic and epigenetic measurements performed on different cells of the same type. MATCHER works by first learning a separate manifold for the trajectory of each kind of genomic data, then aligning the manifolds to infer a shared trajectory in which cells measured using different techniques are directly comparable. Using scM&T-seq data, we confirm that MATCHER accurately predicts true single cell correlations between DNA methylation and gene expression without using known cell correspondence information. We also used MATCHER to infer correlations among gene expression, chromatin accessibility, and histone modifications in single mouse embryonic stem cells. These results reveal the dynamic interplay between epigenetic changes and gene expression underlying the transition from pluripotency to differentiation priming. Our work is a first step toward a united picture of heterogeneous transcriptomic and epigenetic states in single cells.

## Introduction

Understanding the mechanisms that regulate gene expression across space and time is a fundamental challenge in biology. Epigenetic modifications such as DNA methylation, histone marks, and chromatin accessibility are known to regulate gene expression, but the precise details of this regulation are not well understood. Single cell genomic technologies reveal heterogeneity within populations of cells, including complex tissues, tumors, and cells undergoing temporal changes [1, 2]. Furthermore, because bulk data consist of measurements averaged across a population of cells, single cell genomic data enable, in principle, much more precise study of how epigenetic changes and gene expression vary together.

Single cell RNA-seq has been applied with great success to the study of sequential cellular processes such as differentiation and reprogramming [3–7]. In such experiments, each sequenced cell is assumed to be at one point in the process, and sequencing enough cells can reveal the progression of gene expression changes that occur during the process [8, 9]. More recently, several experimental techniques for performing single cell epigenetic have been developed [10–17], and initial analyses have demonstrated that single cell epigenetic data can be used to elucidate the series of changes in a sequential process [16, 18, 19].

Identifying correlations among the epigenome and transcriptome dynamics would allow more complete understanding of the sequential changes that cells undergo during biological processes. Measuring multiple genomic quantities from a single cell, or multi-omic profiling [20, 21], would be the best way to identify such correlations. Unfortunately, performing single cell multi-omic profiling is very difficult experimentally, because an assay on chromatin or RNA destroys the respective molecules and only tiny amounts of DNA and RNA are present in a single cell. In certain cases, it is possible to assay RNA and DNA [14, 22–24] or RNA and proteins [25, 26] from the same single cell, but experimentally performing multiple assays on either chromatin or RNA from the same cell is currently impossible.

Our knowledge of epigenetic regulation suggests that any large changes in gene expression, such as those that occur during differentiation, are accompanied by epigenetic changes. Therefore, it should be possible, in principle, to infer sequential changes in cellular epigenetic state during a process. Furthermore, if cells undergoing a common process are sequenced using multiple genomic techniques, examining any of the genomic quantities should reveal the same underlying biological process. For example, the main difference among cells undergoing differentiation will be the extent of their differentiation progress, whether you look at the gene expression profiles or the chromatin accessibility profiles of the cells.

We reasoned that this property of single cell data could be used to infer correspondence between different types of genomic data. To infer single cell correspondences, we use a technique called manifold alignment [27, 28]. Intuitively, manifold alignment constructs a low-dimensional representation (manifold) for each of the observed data types, then projects these representations into a common space (alignment) in which measurements of different types are directly comparable. To the best of our knowledge, manifold alignment has never been used in genomics. However, other application areas recognize the technique as a powerful tool for multimodal data fusion, such as retrieving images based on a text description, and multilingual search without direct translation [28].

We refer to our method as MATCHER (Manifold Alignment to CHaracterize Experimental Relationships). Using MATCHER, we identified correlations between transcriptomic and epigenetic changes in single mouse embryonic stem cells as they progressed along a trajectory from pluripotency to a differentiation primed state.

## Results

### Overview of MATCHER

Manifold alignment has previously been used to construct a shared, low-dimensional representation that recapitulates known correspondence information between two different kinds of data [27, 28]. The simplest example of manifold alignment is canonical correlation analysis (CCA), in which linear projections of each space are aligned. Gaussian process latent variable models have also been used to perform manifold alignment by learning completely [29, 30] or partially [29] shared latent representations of high-dimensional, multimodal data. Given a set of images and corresponding text descriptions, manifold alignment can be used to identify a low-dimensional representation that allows the prediction of a caption for a new image. This somewhat analogous to the problem of retrieving a corresponding epigenetic measurement for a given single cell transcriptome. However, in the context of single cell genomic data, correspondence information is not generally available to train a model, because it is impossible in most cases to measure more than one quantity on a single cell. Therefore, we developed a novel approach for manifold alignment without correspondence that leverages the unique aspects of this problem.

We assume that:

1. Single cell genomic data from cells proceeding through a biological process lie along a one-dimensional manifold. Another way of saying this is that the variation among cells can be explained mainly by a single latent variable (“pseudotime”) corresponding to position within the process.
2. Each of the genomic quantities under consideration changes in response to the same underlying process.
3. The biological process is monotonic, meaning that progress occurs only in one direction. Processes that alternate between forward and backward progress or repeat cyclically would violate this assumption.
4. The cells in each experiment are sampled from the same population, process, and cell type.

Given these assumptions, there are only three possible types of differences among the one-dimensional manifold representations of each data type: orientation, scale, and “time warping” (Fig. 1a). We can perform manifold alignment without correspondence information by accounting for these three types of differences. Differences in orientation can occur if the biological process corresponds to increasing manifold coordinates for one type of genomic data but decreasing coordinates for another data type. We can reconcile different orientations by simply reversing the order of one set of manifold coordinates. It is not possible to infer the correct orientation from data, so we use biological prior knowledge to choose the correct orientation for the manifold inferred from each type of data. To address scale differences, we can normalize the manifold coordinates to lie between 0 and 1. Time warping effects can occur if different genomic quantities change at different rates. For example, gene expression changes may occur slowly at the beginning of a process and gradually speed up, while changes in chromatin accessibility may show exactly the opposite trend during the process (Fig. 1a). We account for time warping effects by learning a monotonic warping function for each type of data (see below for details).

**Figure 1:**
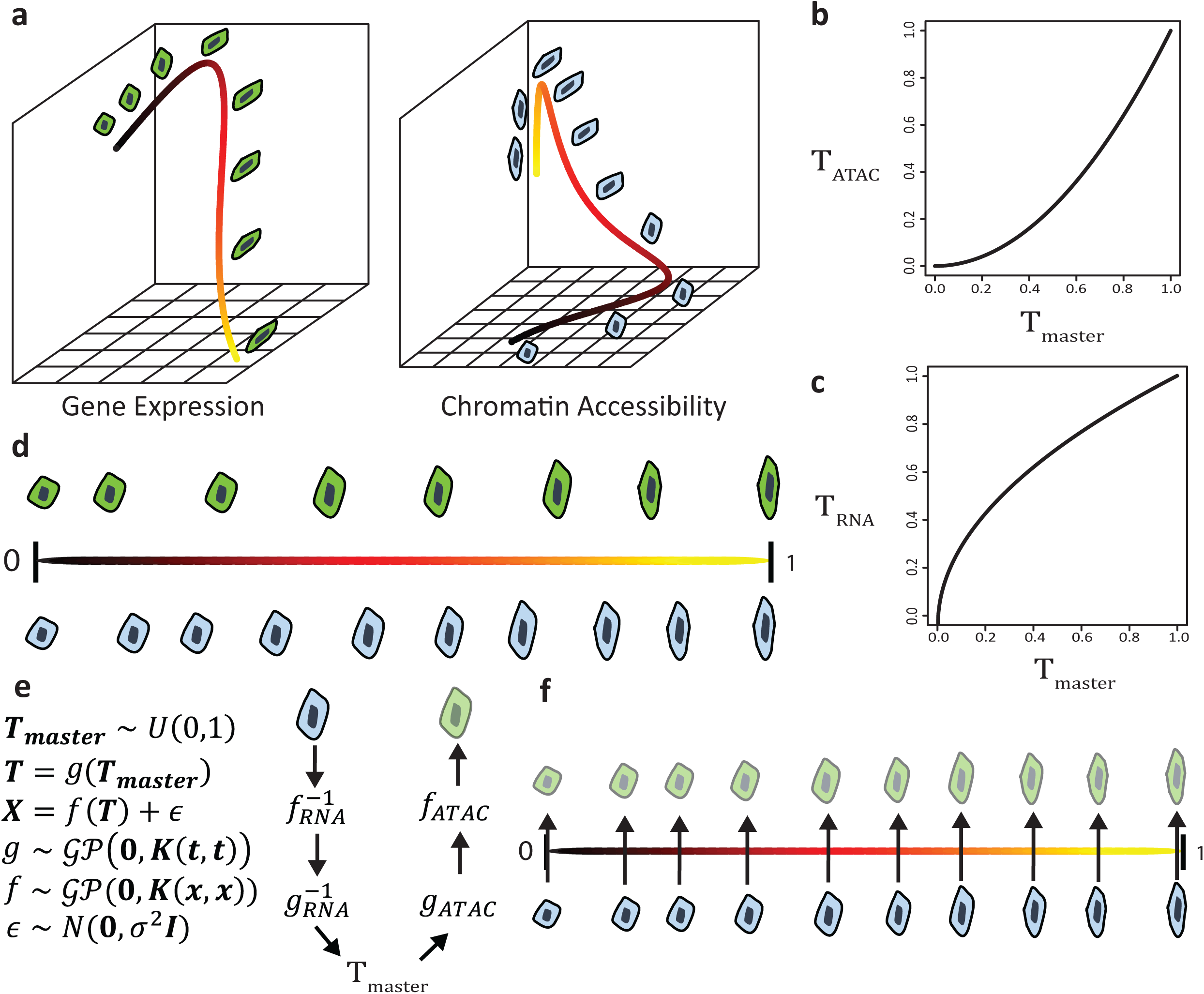
MATCHER Method Overview. (a) We infer manifold representations of each dataset using a Gaussian process latent variable model (GPLVM). However, the resulting “pseudotime” values from different genomic data types are not directly comparable due to differences in orientation, scale, and “time warping”. Both the color of the manifold (black to yellow) and cell morphology (blob to oblong) indicate position within this hypothetical process. (b)-(c) To account for these effects, pseudotime for each kind of data is modeled as a nonlinear function (warping function) of master time using a Gaussian process. (d) MATCHER infers “master time” in which the steps of a biological process correspond to values uniformly distributed between 0 and 1 and are comparable among different data types. However, different datasets are measured from different physical cells, and thus may sample different points in the biological process and even different numbers of cells. (e) Diagram showing how MATCHER’s generative model can infer corresponding cell measurements. The generated cell is drawn with transparency to indicate that this is an inferred rather than observed quantity. (f) Applying MATCHER to multiple types of data provides exactly corresponding measurements from observed cells and unobserved cells (indicated with transparency) generated by MATCHER.

We use a Gaussian process latent variable model (GPLVM) to infer pseudotime values separately for each type of data. A GPLVM is a nonlinear, probabilistic, generative dimensionality reduction technique that models high-dimensional observations as a function of one or more latent variables [31]. The key property of a GPLVM is that the generating function is a Gaussian process, which allows Bayesian inference of latent variables nonlinearly related to the high-dimensional observations [32, 33]. The nonlinear nature of this model makes it more flexible and robust to noise than a linear model such as principal component analysis (PCA). In fact, PCA can be derived as a special case of a GPLVM in which the Gaussian process generating function uses a linear kernel [31]. Importantly, GPLVMs are also generative models, meaning that they can answer the counterfactual question of what an unobserved high-dimensional datapoint at a certain location on a manifold *would* look like. The generative nature of GPLVMs is particularly important to our approach: We use this property to infer correspondence among single cell genomic quantities measured in different ways. We note that GPLVMs have previously been used to infer latent variables underlying differences among single cell gene expression profiles [34–36]; our approach differs from these previous approaches in that we use GPLVMs as part of a *manifold alignment* approach and *generate* measurements from unobserved cells to *integrate* multiple types of single cell measurements.

After inferring pseudotime separately for each type of data, we learn a monotonic warping function (Fig. 1b-c) that maps pseudotime values to “master time” values, which are uniformly distributed between 0 and 1 (Fig. 1d). This is equivalent to aligning the quantiles of the pseudotime distribution to match the quantiles of a uniform random variable. Master time values inferred from different data types are then directly comparable, corresponding to the same points in the underlying biological process.

The model (Fig. 1e) that we use to infer master time values allows us to *generate* corresponding cell measurements even from datasets where the measurements were performed on different single cells. The different types of measurements may produce datasets with cells from different positions in the biological process, and even different numbers of cells (Fig. 1e). To generate a corresponding measurement for a cell, we take the master time value inferred for a given cell, such as one measured with RNA-seq. Then we map this master time value through the warping function to a pseudotime value for a different type of data, such as ATAC-seq. Using the GPLVM trained on ATAC-seq data, we can output a corresponding cell based on this pseudotime value. As Fig. 1f shows, the generative nature of the model allows MATCHER to infer what unobserved cells measured with one experimental technique *would* look like if they corresponded exactly to the cells measured using a different technique. These corresponding cell measurements can then be used in a variety of ways, such as computing correlation between gene expression and chromatin accessibility.

Although it is very difficult in general to measure multiple genomic quantities on the same single cell, one particular protocol (scM&T-seq) has been developed for measuring DNA methylation and gene expression in the same single cell [14]. It is possible that future protocols will enable other joint measurements. In such cases, we can incorporate the known correspondence information into our model by using a shared GPLVM [29] to infer a shared pseudotime latent variable for both data types.

### Data Description and Processing

Several high-throughput single cell versions of epigenetic assays have been developed, including single cell bisulfite sequencing (DNA methylation) [14], ATAC-seq (chromatin accessibility) [13], and ChIP-seq (histone modification) [12]. Each of the initial studies that pioneered these methods applied them to mouse embryonic stem cells (mESCs) grown in serum, a classic model system of stem cell biology. Cells in this system are heterogeneous, differing depending on where they are located along a spectrum ranging from a pluripotent ground state to a differentiation primed state [37]. Note that mESCs grown in serum have different properties than mESCs cultured in 2i medium, which are much more homogeneous and differ primarily in their cell cycle stage [34, 37].

We collected the publicly available data from these papers. In total, we have four kinds of single cell data from a total of 4,974 cells: 250 cells with gene expression data [37], 61 with DNA methylation [14], 76 with chromatin accessibility [13], and 4,587 with H3K4me2 ChIP [12]. The DNA methylation data were collected using the scM&T-seq protocol, which measures gene expression and DNA methylation simultaneously in a single cell [14].

The processing of single cell epigenetic data is more difficult than RNA-seq, because the epigenetic data are nearly binary at each genomic position (apart from allele-specific effects and copy number variations) and extremely sparse, with only a few thousand reads per cell in many cases. This makes it very difficult to extract any meaningful information at base pair resolution from a single cell. Instead, we followed the data processing steps laid out in each of the respective papers that developed these techniques and aggregated the reads across related genomic intervals. For example, we followed the authors’ lead in summing the chromatin accessibility data values from ATAC-seq in a given cell across all of the binding sites for a given transcription factor. Doing this for each of 186 transcription factors results in a matrix of 186 chromatin accessibility signatures across the set of cells. The DNA methylation data and H3K4me2 ChIP-seq data were aggregated in a similar way. We obtained the processed DNA methylation and ChIP-seq data from the initial publications. The processed ATAC-seq data are not publicly available, so we processed the data by implementing ourselves the pipeline described in the paper. We found that the DNA methylation data showed the highest detection rate per cell; the ChIP-seq data had the lowest detection rate. Consequently, we were able to aggregate the DNA methylation data over relatively small genomic intervals such as individual promoters or CpG islands.

### Single cell transcriptome and epigenome data show common modes of variation

It seems likely that gene expression, DNA methylation, chromatin accessibility, and histone modifications will all change during the transition from pluripotency to a differentiation primed state. However, we wanted to investigate that this crucial assumption holds in this particular system.

To test our hypothesis that each of these epigenetic data types are changing over the course of a common underlying process, we first attempted to construct a cell trajectory for each type of data. Using SLICER, a method we previously developed [9], we visualized each type of data as a two-dimensional projection and inferred a one-dimensional ordering for the cells. The 2D projections show that each type of data resembles a one-dimensional trajectory rather than a 2D blob of points (Fig. 2a-d). Note that these 2D projections do not force the data into a one-dimensional shape; the plots could look like a diffuse point cloud, and the fact that they instead resemble trajectories shows that the differences among cells are predominantly one dimensional. Furthermore, the projections of each kind of data are strikingly similar visually (Fig. 2a-d).

**Figure 2:**
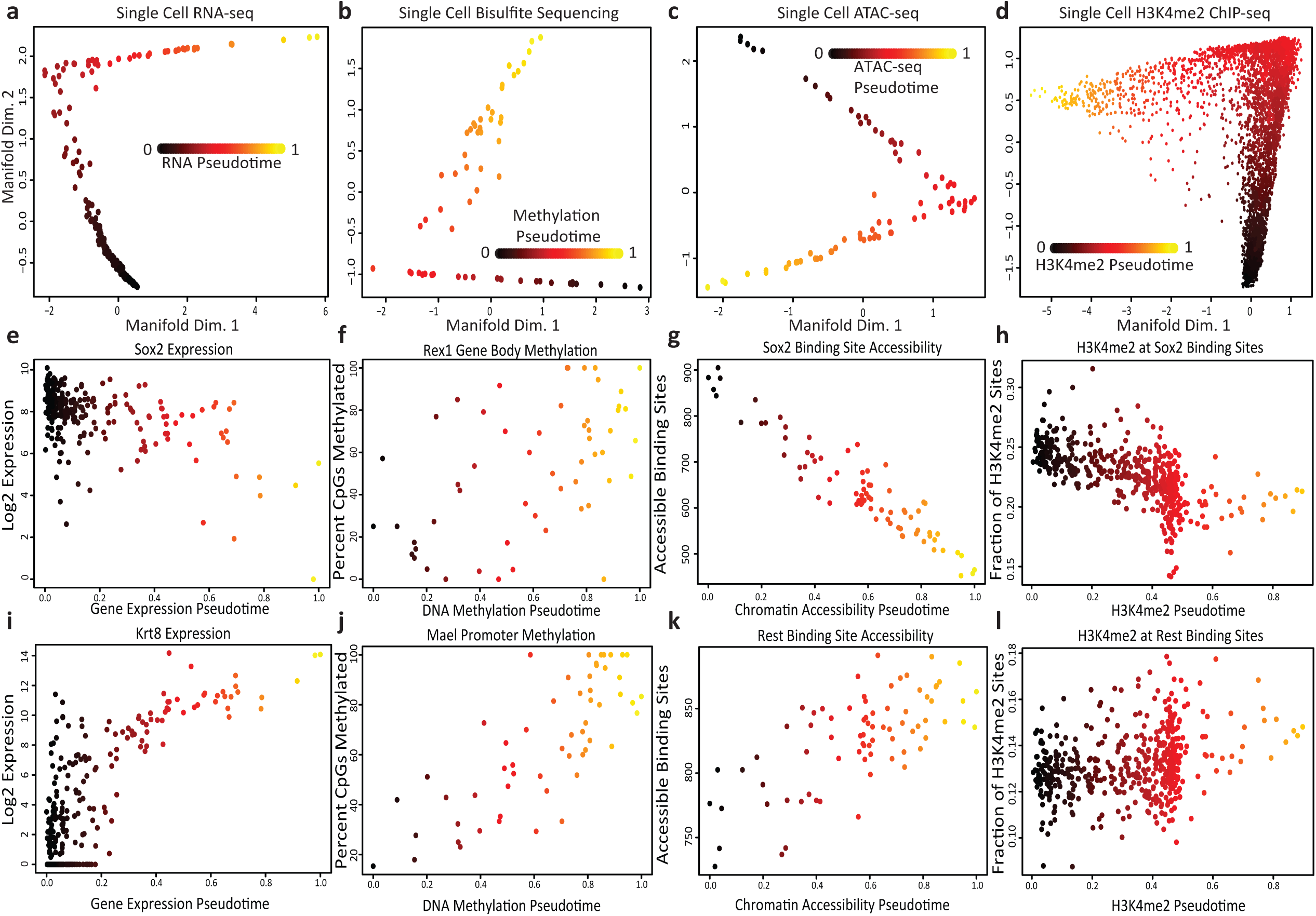
Single cell transcriptome and epigenome data show common modes of variation. (a)-(d): Single cell trajectories constructed by SLICER from RNA-seq, bisulfite sequencing, ATAC-seq, and H3K4me2 ChIP-seq of mouse embryonic stem cells grown in serum. (e)-(l) Levels of important gene expression, DNA methylation, chromatin accessibility, and H3K4me2 markers across the trajectories. Note: We used SLICER for the analysis in this figure because it is a previously published method for constructing cell trajectories that allowed us to investigate the hypothesis that single cell transcriptome and epigenome measurements share common sources of variation. SLICER and MATCHER are completely separate methods; MATCHER does not rely on SLICER in any way; and SLICER could not be used to integrate multiple types of measurements as MATCHER does, because SLICER lacks the ability to generate unobserved cell measurements.

We further investigated these trajectories to determine whether they correspond to the same underlying process. The trajectory built from RNA data shows decreasing expression of pluripotency genes such as SOX2, consistent with previously published analyses [37] (Fig. 2e). DNA methylation of the gene body of *Rex1*, a gene that is shut off during the transition from pluripotency to differentiation priming [38], increases during the process (Fig. 2f). The single cell ATAC-seq data show that the chromatin accessibility of binding sites for the SOX2 transcription factor decreases over pseudotime (Fig. 2g). Similarly, the levels of H3K4me2, a histone modification associated with active enhancers and promoters, decrease at SOX2 binding sites (Fig. 2h). The RNA-seq data show increasing expression of previously identified differentiation markers [37] such as *Krt8* (Fig. 2i). DNA methylation of the promoter for *Mael* increases, consistent with previous findings [38] (Fig. 2j). Both the chromatin accessibility (Fig. 2k) and H3K4me2 levels (Fig. 2l) at REST binding sites increase, consistent with the known role of REST in repressing key lineage-specifying genes [39, 40]. In summary, our analysis indicates that each type of single cell data varies along a trajectory, establishing a continuum that ranges from pluripotency to a differentiation primed state.

We used SLICER to perform this initial exploratory analysis, but for the rest of this study, we use MATCHER, which is completely separate from SLICER and does not rely on the method in any way. We did confirm, however, that the master time values inferred by MATCHER are highly correlated with the pseudotime values inferred by SLICER (Supplementary Figure 1). Note also that SLICER cannot be used to integrate multiple types of single cell measurements in the way the MATCHER does, because the model underlying SLICER is not generative.

### MATCHER accurately models synthetic and real data

To evaluate the accuracy of MATCHER, we generated synthetic data for which ground truth master time is known. We generated data by sampling 100 master time values uniformly at random from the interval [0,1], then mapping these to pseudotime values through a warping function. Using the resulting pseudotime values, we generated 600 “genes” each following a slightly different “expression pattern” (function of pseudotime). Normally distributed noise was added to each gene expression value. We then used MATCHER to infer master time from these simulated gene expression values, and measured accuracy as the correlation between true and inferred master time values. Note that we use Pearson rather than Spearman correlation because we expect true and inferred master time to be linearly related (equal, in fact), and a nonlinear relationship would indicate that the inference process is inaccurate. The results of our simulations indicate that MATCHER accurately infers master time across a range of different warping functions and noise levels (Supplementary Figs. 2-3). The method is very robust to noise in the simulated genes, yielding a correlation of 0.92 at a noise level of σ = 9, which is greater than 50% of the range of the simulated features.

We also tested MATCHER on real data. We used scM&T-seq data, in which DNA methylation and gene expression are measured in the same single cells [14], so that the true correspondence between single cell measurements is known. Note that we used the known cell correspondence information for validation only, not during the inference process. We first checked the relationship between master time inferred by MATCHER from RNA-seq and DNA methylation data by calculating the correlation between inferred master time values for corresponding DNA methylation and RNA-seq cells. This showed that the master time values, although not identical, are highly concordant (Pearson *ρ* = 0.63). Predicting covariance of multiple genomic quantities across single cells is one of the key applications of MATCHER. Therefore, as an additional test, we investigated whether MATCHER can accurately infer correlations between DNA methylation events and gene expression. Here, we used Spearman correlation because we are interested in both linear and nonlinear relationships. We selected a set of genes and proximal methylated loci that showed statistically significant correlation in the original analysis of the scM&T-seq data [14]. Angermueller et al. grouped these pairs according to the type of region where the methylation site occurred. We selected the three types of regions with the largest number of significant pairs (low methylation regions, H3K27me3 peaks, and P300 binding sites). Then, for each significant pair, we compared the true correlation (calculated using true cell correspondences) and correlation inferred by MATCHER (calculated using inferred cell correspondences). We also used MATCHER to compute correlations for the same gene-locus pairs using a single cell RNA-seq dataset published by a different lab [37]. In this dataset, the cells measured using RNA-seq are the same cell type, but not the same physical cells as those assayed for DNA methylation by Angermueller et al. In both cases, the inferred correlations closely match the true correlations (Fig. 3). The correlations computed using the Kolodziejczyk data show slightly less concordance with the ground truth, likely due to the inevitable biological and technical variation that occur when different labs repeat an experiment. Even so, the vast majority of inferred correlations have the correct sign, and the relative magnitude of correlations tends to be preserved.

**Figure 3:**
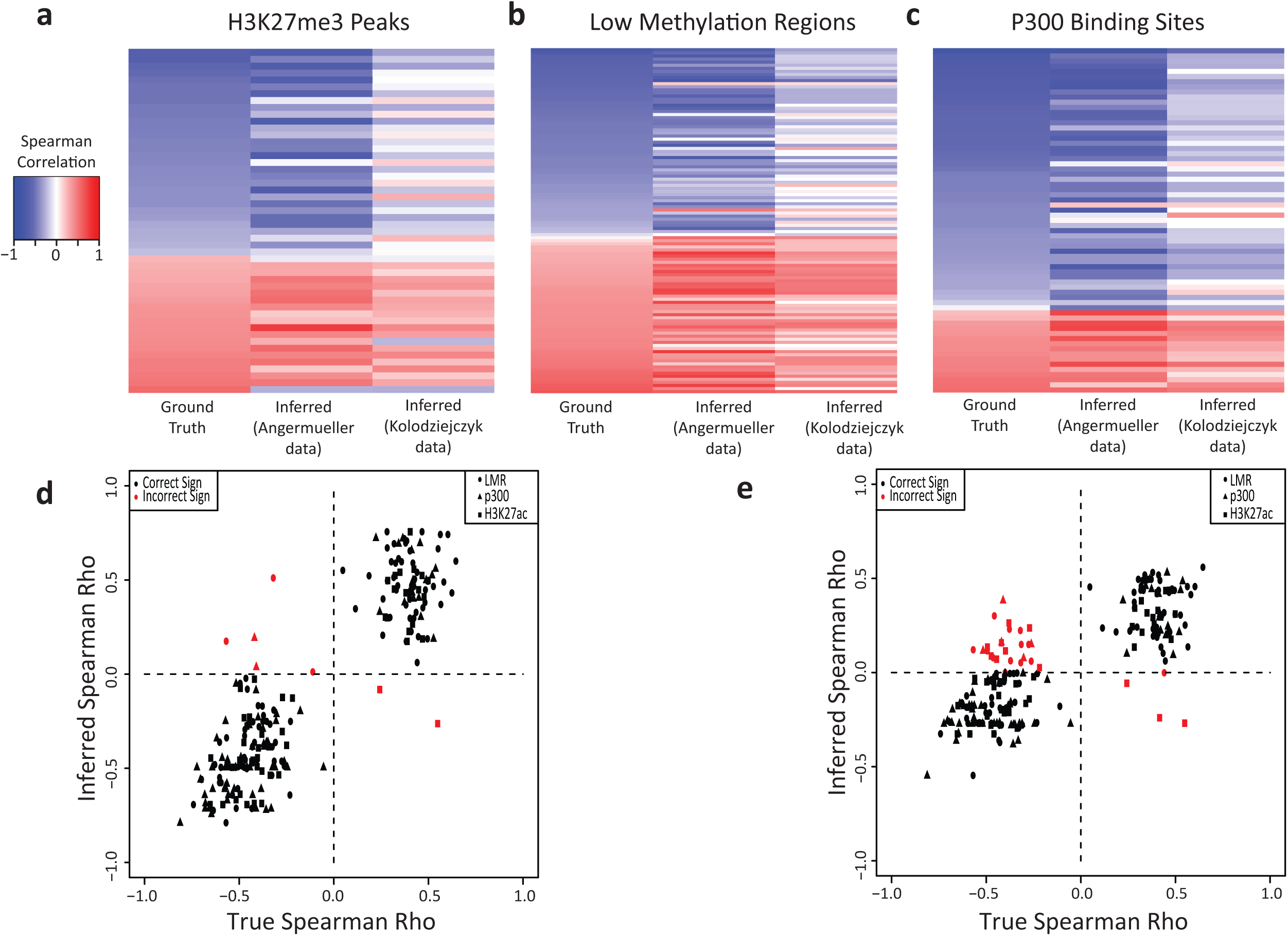
MATCHER accurately infers known correlations between DNA methylation and gene expression. (a)-(c) Heatmaps comparing true correlations between gene expression and DNA methylation of related regions (H3K27me3 peaks, LMRs, and P300 binding sites). The first column of each heatmap shows the true correlation based on known correspondence information, the second column shows the correlation inferred by MATCHER in the same dataset, and the third column is correlation inferred by MATCHER using a completely different single cell RNA-seq dataset from mESCs grown in serum. (d)-(e) Scatterplot representation of the results shown in (a)-(c). Panel (d) contains correlations computed using the Angermueller data; panel (e) is correlations computed from the Kolodziejczyk data. Each point represents the true and inferred correlation for a single gene-site pair; ideal results would lie along the *y* = *x* line. Note that the sign of the inferred correlation is correct for the vast majority of pairs.

### Correlations among single cell gene expression, chromatin accessibility, and histone modifications

We next used MATCHER to investigate the relationships among gene expression, chromatin accessibility, and histone modifications during the transition from pluripotency to a differentiation primed state. To our knowledge, this is the first time that investigation of the relationship among these three genomic quantities has been performed in single cells.

Because H3K4me2 is a histone modification associated with promoter and enhancer activation, we expect levels of the modification to correlate positively with chromatin accessibility. We confirmed this is, indeed, the case by inferring correlations between chromatin accessibility and H3K4me2 at the respective regions bound by 186 transcription factors and DNA binding proteins (Fig. 4a). The majority of these correlations are strongly positive, indicating that activating histone modifications and chromatin accessibilty tend to change in unison during preparation for differentiation.

**Figure 4:**
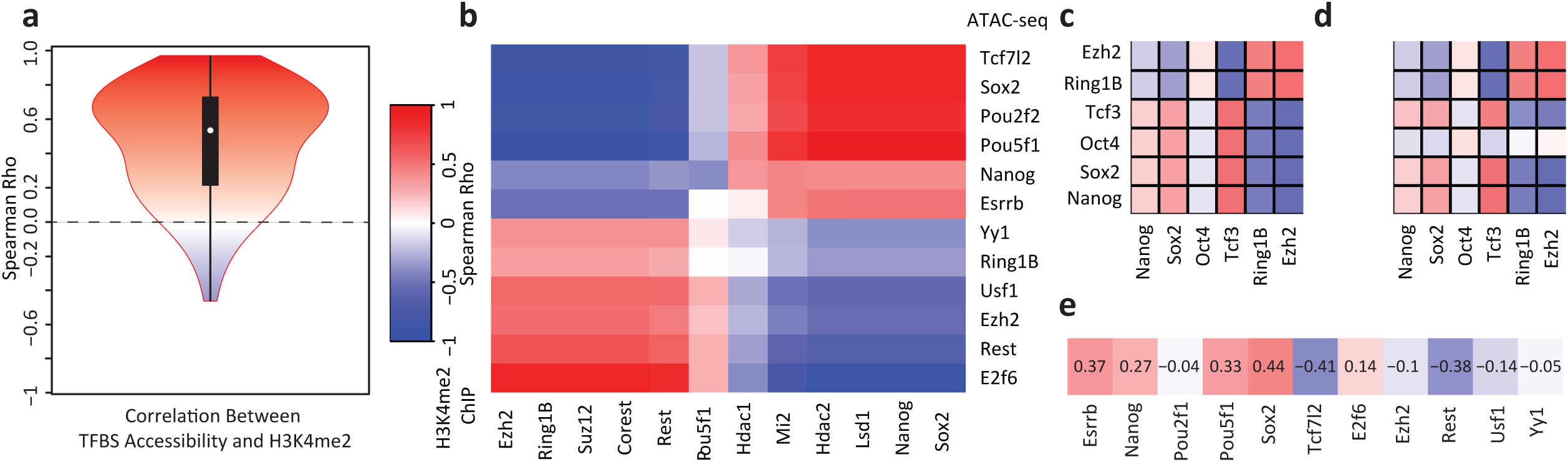
Correlations among single cell gene expression, chromatin accessibility, and histone modifications. (a) Violin plot of correlations among chromatin accessibility and H3K4me2 of transcription factor binding sites for 186 transcription factors. Note that most correlations are strongly positive. (b) Correlation between chromatin accessibility and H3K4me2 data reveals that targets of pluripotency factors/NuRD complex and targets of Polycomb Group/Trithorax Group proteins are anticorrelated in single cells. (c) Correlation between gene expression signatures and chromatin accessibility signatures. (d) Correlation between gene expression signatures and H3K4me2 signatures. (e) Correlation between gene expression of DNA binding proteins and chromatin accessibility of their targets.

While investigating the correlation between H3K4me2 and chromatin accessibility, we found that the genomic binding regions clustered into two main groups: (1) pluripotency transcription factors and the NuRD complex and (2) chromatin remodeling factors that repress or activate lineage specific genes (Fig. 4b). Rotem et al. noted a similar relationship in the H3K4me2 data [12]. The accessibility of binding sites for OCT4 (also known as POU5F1), NANOG, and SOX2, well-established pluripotency transcription factors, is strongly anticorrelated with the accessibility of binding sites for EZH2, RING1B, and SUZ12, which are Polycomb Group proteins (PcG) [41]. The targets of the transcription factor YY1, which recruits PcG proteins [42], show a similar trend to the PcG proteins. Given that PcG proteins play a key role in repressing neuronal lineage genes in pluripotent cells [43], this anticorrelation suggests that chromatin is being remodeled to prime lineage-specific genes while shutting down regions associated with pluripotency. REST and COREST show a similar pattern to the PcG proteins; these proteins are known to co-associate with the polycomb repressive complex (PRC2) and also to repress key lineage specific genes in pluripotent cells [39, 40]. Interestingly, the targets of USF1, which is known to recruit Trithorax Group (TrxG) proteins [44], also show a pattern of increasing chromatin accessibility. The TrxG proteins are chromatin activators that regulate lineage differentiation genes [43–45], suggesting that the activation of certain differentiation genes is occurring while their repression by PRC2 is being lifted. Finally, targets of LSD1, MI2, HDAC1, and HDAC2, components of the NuRD complex, show positive correlation with targets of pluripotency factors. The NuRD complex contains chromatin remodeling proteins that remove histone methylation and histone acetylation marks and function to “decommission” pluripotency enhancers during early differentiation [46]. In summary, our analysis of correlation between chromatin accessibility and H3K4me2 marks indicate that the overall trend in both types of data is toward chromatin changes that shut off pluripotency and begin to lift lineage repression in preparation for differentiation.

We also computed correlations between gene expression and chromatin accessibility, and between gene expression and H3K4me2. To identify populations of RNA molecules with a clear relationship to the aggregated genomic regions used to compute chromatin accessibility measurements, we combined gene expression levels from genes whose promoters overlapped the binding sites for several proteins. We looked specifically at binding regions for EZH2, RING1B, TCF3, OCT4, SOX2, and NANOG. After locating genes whose promoters overlapped each of these binding regions, we filtered the sets of genes to remove genes that occurred in multiple binding regions. We then normalized the expression of each gene (zero mean, unit variance) and calculated the aggregate expression for each set of genes. The aggregate expression of these sets of genes correlates well with the chromatin accessibility and H3K4me2 of the gene promoters (Fig. 4c-d), with the exception of OCT4. The expression of OCT4 targets are only weakly correlated with the aggregate chromatin accessibility and H3K4me2. Supplementary Figures 4 and 5 show the corresponding values inferred by MATCHER for gene expression, chromatin accessibility, and H3K4me2 values in the same single cells.

Finally, we asked how the RNA expression levels of key pluripotency factors and chromatin remodeling proteins relate to the chromatin accessibility of their binding sites (Fig. 4e). Using the same transcription factors and DNA binding proteins as in Fig. 4a, we calculated the correlation between the expression level of each gene and the overall chromatin accessibility of the sites where its protein product binds to the genome. The pluripotency transcription factors ESRRB, NANOG, POU5F1, and SOX2 each show positive correlation between expression and chromatin accessibility. This indicates that the expression of these genes is being shut off at the RNA level at the same time as the binding of the factors is shut off at the chromatin level. Interestingly, *Tcf7l2* expression shows strong negative correlation with the chromatin accessibility of its targets. We speculate that this negative correlation may be due to the fact that TCF7L2 functions primarily as a transcriptional repressor [47], and thus increased expression will lead to more repression of its targets. In contrast to the pluripotency factors, the expression of genes involved in chromatin remodeling show weak negative correlation with the accessibility of their binding sites. The fact that these correlations are nearly zero indicates that changes in the chromatin accessibility of the targets of these chromatin remodeling complexes occurs primarily without accompanying changes in the gene expression levels of the remodelers. The one exception is *Rest*, whose expression shows strong negative correlation with the accessibility of its binding sites.

### Warping functions inferred by MATCHER suggest rapid transition between two metastable states

Upon inspecting the warping functions inferred by MATCHER for the gene expression, chromatin accessibility, and H3K4me2 data, we noticed that the curves all had similar shapes (Fig. 5a-c). The Angermueller single cell RNA-seq dataset [14] also shows a similar pattern (see Supplementary Figure 6 for plots of all warping functions). The warping functions are nearly flat at the beginning and end of pseudotime, and steeply sloped in between. One possible explanation for this pattern is a process in which cells transition rapidly from one metastable state to another. We hypothesize that the shapes of the warping functions may reflect the biology of embryonic stem cells grown in serum, in which some pluripotent cells begin to “lose control” and transition to a differentiation primed state [37, 38]. In support of this hypothesis, a recent paper utilizing single cell FISH characterized such a transition between metastable states in mouse embryonic stem cells [38]. Interestingly, the warping function for the DNA methylation data does not show this switch-like behavior (Fig. 5d). We suspect that this may be due to partial decoupling of DNA methylation and gene expression changes (see next section).

**Figure 5:**
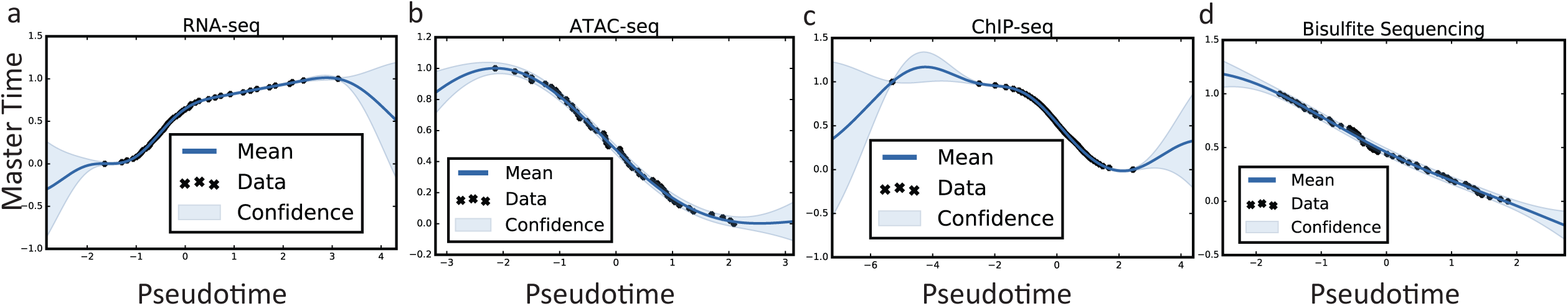
Warping functions inferred by MATCHER suggest rapid transition between two metastable states. (a)-(d) Gaussian process warping functions for (a) RNA-seq, (b) ATAC-seq, (c) <u>ChIP</u>-seq, and (d) <u>scM</u>&T-seq. Note: Because the mean of Gaussian process warping function for <u>ChIP</u>-seq is not monotonic over the observed data range, we used linear interpolation for the <u>ChIP</u>-seq warping function in all of the analyses reported in the paper.

### Incorporating known cell correspondence information

The study that pioneered scM&T-seq found variability in the strength of coupling between gene expression and DNA methylation across the set of cells [14]. We investigated this phenomenon further by plotting DNA methylation master time as a function of RNA master time (Fig. 6a). This plot revealed an intriguing trend: DNA methylation and RNA master time track together quite well until a specific point in RNA master time. After that point, the degree of coupling suddenly decreases. We speculate that this trend may occur because specific DNA methylation changes are required to trigger a key step in the process of gene expression changes that occur moving from a pluripotent to a primed state. After this point in the process, the sequential gene expression changes proceed somewhat independently from the DNA methylation changes. In support of this hypothesis, the cells in which DNA methylation and gene expression match show high levels of *Rex1* expression, while the remaining cells show much lower expression (Fig. 6a). The *Rex1* gene was previously shown to be a marker for two distinct metastable expression states in mouse embryonic stem cells [38]. The transition between these two states is dependent on the activity of DNA methyltransferase (DNMT) enzymes, and knocking out DNMT activity greatly increases the proportion of cells in the *Rex1*-high state [38].

**Figure 6:**
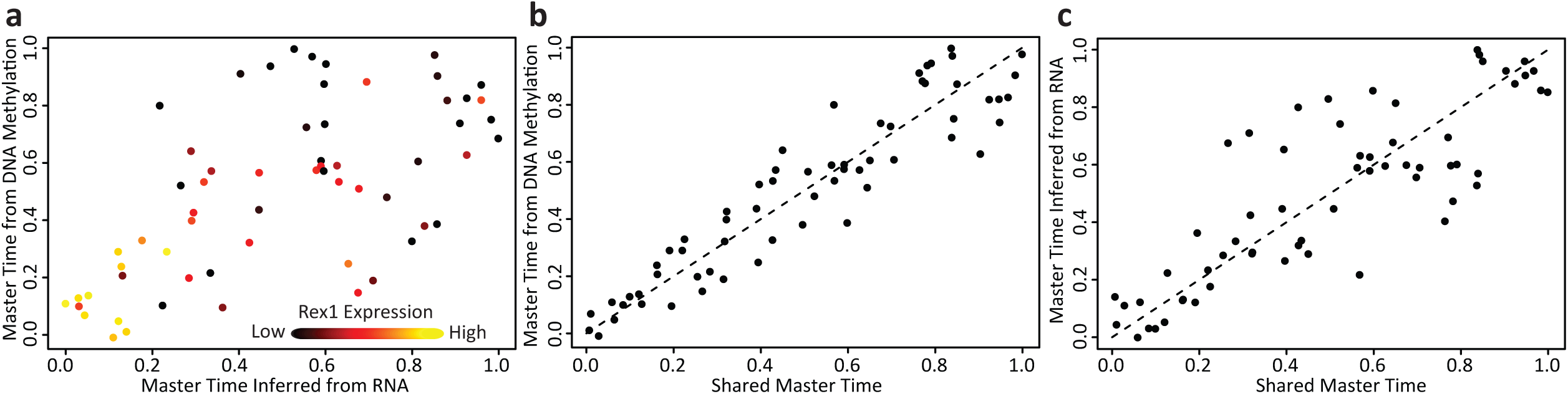
Incorporating known cell correspondence information. (a) Scatterplot of master time inferred using gene expression (x-axis) and DNA methylation (y-axis) from the same single cells. Points are colored by the expression of *Rex1*. (b) Scatterplot of shared master time inferred from both gene expression and DNA methylation (x-axis) and master time inferred using DNA methylation only (y-axis). (c) Scatterplot of shared master time inferred from both gene expression and DNA methylation (x-axis) and master time inferred using gene expression only (y-axis).

To test the significance of the partial decoupling between DNA methylation and gene expression, we computed separate Pearson correlation values for cells with RNA-seq master time less than 0.5 and greater than 0.5. Then we performed Fisher’s *r*-to-*z* transformation on the correlations and computed a *p*-value for the null hypothesis that the two correlations are equal (one-tailed test). The *p*-value was 0.0039, indicating a highly significant difference.

We next used MATCHER to infer a shared master time value using both DNA methylation and gene expression data for each cell (Fig. 6b-c). The resulting shared master time values reconcile the sequence of changes occurring in both genomic quantities. The Pearson correlation between DNA methylation master time and RNA master time is 0.63. In contrast, the correlation between DNA methylation master time and shared master time is 0.93, and the correlation between RNA master time and shared master time is 0.84. Thus, we have used MATCHER to infer master time that reflects the known correspondence information available from scM&T-seq data. This demonstrates the capability of MATCHER to model single cell data either with or without the use of known correspondence information.

## Discussion

In this study, we used MATCHER to characterize the corresponding transcriptomic and epigenetic changes in embryonic stem cells undergoing the transition from pluripotency to a differentiation primed state. Interesting future directions of research include extending the model to align manifolds with dimensionality higher than one, as well as adapting the method for cell populations whose cells fall into discrete clusters rather than along one continuous spectrum. In addition, our model does not explicitly account for branching trajectories, which can arise in biological processes with multiple outcomes [3, 9]. A simple way to handle such situations would be to assign cells to branches before running MATCHER, and then perform manifold alignment on each branch separately.

Although the Hi-C protocol for measuring chromatin conformation has been adapted to single cells [10], we did not include single cell Hi-C data in this study for two reasons. First, to the best of our knowledge, there are no published single cell Hi-C datasets from mouse embryonic stem cells. In addition, Hi-C data are a set of pairwise interactions (a matrix for each cell, rather than a vector), and it is not clear how to construct a trajectory from this type of data. Further work is necessary to investigate whether chromatin conformation shows sequential changes during biological processes, as well as the best ways infer such sequential changes and integrate them with other types of data.

One promising application of the method is aggregating single cell measurements into biologically meaningful groups. Cells can be grouped by their inferred master time values, and measurements within these groups can be aggregated. In experiments with thousands of cells, this will likely enable correlation between individual loci and related genes, which is currently impossible because of the extreme sparsity of the epigenetic data. Computational aggregation of measurements from many similar single cells may be the most immediate way to address the sparsity of single cell epigenetic measurements, although experimental protocols will likely improve over the long term.

MATCHER gives insight into the sequential changes of genomic information, allowing the use of both single cell gene expression and epigenetic data in the construction of cell trajectories. In addition, it reveals the connections among these changes, enabling investigation of gene regulatory mechanisms at single cell resolution. MATCHER promises to be useful for studying a variety of biological processes, such as differentiation, reprogramming, immune cell activation, and tumorigenesis.

## Methods

### RNA-seq Data Processing

We obtained the processed RNA-seq data for 250 cells from Kolodziejczyk et al. [37] In the original paper, gene quantification was performed using read counts that were normalized for sequencing depth and batch effects [37]. We log transformed these normalized counts and used our previously published neighborhood variance method to select an informative subset of genes to feed into MATCHER.

To identify populations of RNA molecules with a clear relationship to the aggregated genomic regions used to compute chromatin accessibility and histone modification measurements (see below), we computed analogous aggregated gene expression measurements. We did this by identifying genes whose promoters overlap binding sites for each of 6 proteins (EZH2, RING1B, TCF3, OCT4, SOX2, and NANOG). We then filtered the gene lists so that a given gene appears on only one of the six lists. Then we scaled and centered each gene to have zero mean and unit variance and computed the sum of the genes on each list per cell, as well as the total sum of expressed genes in each cell. The final values used to compute correlations shown in Fig. 4c-d are the centered and scaled differences of the sum for each list of genes and the total sum of gene expression per cell.

### ATAC-seq Data Processing

The processed single cell ATAC-seq data are not publicly available, so we implemented the data processing pipeline described by Buenrostro et al. [13] For each cell, we aligned reads to mm10 using bowtie2, removed PCR duplicates, and counted the number of reads aligning to each of the 50000 peaks identified in the initial paper [13]. We converted these integer read counts, which are predominantly 1 or 0 at a given peak, into binary values (1 for accessible chromatin, 0 for inaccessible) to avoid potential confounding factors that could cause high counts such as copy number variations and repeat elements. Then we used FIMO [48] to identify, for each peak, which of the 186 transcription factor motifs in the JASPAR database [49] occurs in the peak region. Using this peak-to-TF mapping, we aggregated the peak counts for each cell by summing the peaks for each transcription factor motif. This gave a matrix with 186 features across 96 cells. We subsequently removed all cells with fewer than 1000 peaks detected per cell, leaving 77 cells. Dimensionality reduction using PCA and a GPLVM on the 77 cells indicated that one cell was a significant outlier, so we removed this additional cell. The remaining 76 cells were used for all subsequent analyses. We then normalized the 186 × 76 count matrix to account for differences among cells in numbers of peaks detected. We normalized the value of *f*_*ij*_ (feature *i* in cell *j*) by multiplying by the following scale factor: 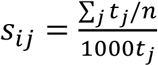, where *t*_*j*_ is the total number of accessible peaks in cell *j*. (The 1000 in the denominator of the scale factor scales the measurements so that the *f*_*ij*_ are close to 1.)

### ChIP-seq Data Processing

We obtained the processed data from Rotem et al. [12], which consists of H3K4me2 ChIP-seq reads from 4587 cells, aggregated using 91 chromatin signatures. We found that these data required further normalization for the total sum of signature values per cell. We normalized the value of *f*_*ij*_ (signature *i* in cell *j*) by multiplying by the following scale factor: 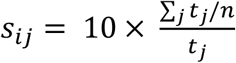, where *t_j_* is the total sum of signatures in cell *j*. (The 10 in the numerator of the scale factor scales the measurements so that the *f*_*ij*_ are close to 1.)

### scM&T-seq Data Processing

RNA-seq and DNA methylation data from Angermueller et al. [14] are publicly available in fully processed form, so we did not perform any further processing. In the original paper, the gene expression levels were computed by counting unique molecular identifiers (UMIs) and subsequently normalized. The DNA methylation values from Angermueller were also normalized in the original paper [14].

We initially tried using the methylation values from all positions in the genome, but PCA and GPLVM results on the full dataset showed no systematic variation related to pluripotency and differentiation. This is likely because only a subset of methylation sites show systematic biological variation in excess of technical variation during the transition from pluripotency to differentiation priming. We therefore selected methylation sites based on a previously validated marker, *Mael*, whose methylation is known to change during the transition to a differentiation primed state [38]. We selected all methylation sites whose correlation with the promoter methylation of *Mael* was at least 0.2. This gave a set of approximately 13,000 methylation sites. There were essentially no methylation sites anticorrelated with *Mael*, consistent with the fact that pluripotent cells are globally demethylated, so that methylation changes in preparation for differentiation occur primarily in a single direction. We also found that using only data from low methylation regions (LMRs), which are known to change methylation state dramatically during differentiation, gives similar results [50].

### Inferring Pseudotime and Learning Warping Functions

We infer pseudotime using a Gaussian process latent variable model (GPLVM) with a single latent variable ***t***. For a more thorough introduction to Gaussian processes and GPLVMs, see Rasmussen [51] or Damianou [33]. Under our model, the observed high-dimensional data (RNA-seq, ATAC-seq, ChIP-seq, DNA methylation, etc.) are generated from ***t*** by a function *f* with the addition of Gaussian noise:

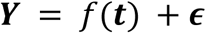

where *∊* ∼ 𝒩 (**0**, σ^2^***I***).

The key property of a GPLVM is that the prior distribution of *f* is a Gaussian process:

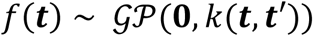

A linear kernel yields a model equivalent to probabilistic PCA, but if we choose the kernel function *k* to be nonlinear, the GPLVM can infer nonlinear relationships between *t* and *Y*. We use the popular radial basis function (RBF) kernel, also called the squared exponential kernel.

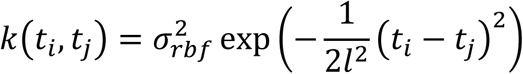

Because a Gaussian process is a collection of random variables for which the covariance of any finite set is a multivariate Gaussian, we have:

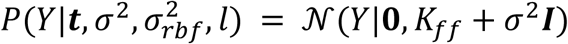

where *K*_*ff*_ is the covariance matrix defined by the kernel function *k*. A simple approach to inferring the latent variable ***t*** would be to find the values that maximize the posterior distribution:

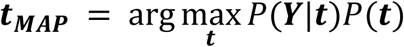

Instead of MAP estimation, we use the method of Damianou [33], which estimates the posterior using a variational approximation. A key advantage of this approach is that it provides a distributional estimate of the latent variables rather than just a point estimate. The approximation relies on the introduction of auxiliary variables called inducing inputs to derive an analytical lower bound on the marginal likelihood. Inference is then performed by maximizing the lower bound with respect to the inducing inputs and the hyperparameters 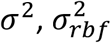, and *l*. We used 10 inducing inputs for all of our analyses, although we confirmed that the results are robust to the number of inducing inputs used.

To learn warping functions from pseudotime to master time, we compute the sample quantiles of pseudotime for a specified number of quantiles, then align these sample quantiles with the theoretical quantiles of a uniform (0,1) random variable. We used 50 quantiles for all analyses in the manuscript, but found that the warping functions are robust to the number of quantiles used. Gaussian process regression is an attractive choice for learning a warping function due to the capability to capture nonlinear effects and uncertainty, but Gaussian processes are not theoretically guaranteed to be monotonic. In practice, we found that the mean of the Gaussian process fit is monotonic in most cases, because the training data are monotonically increasing quantiles. For cases when the mean of the Gaussian process is not monotonic (as is the case for the single cell ChIP-seq data), we use linear interpolation. The monotonicity of the quantiles guarantees that the linear interpolation will be monotonic.

## Figure Legends

**Supplementary Figure 1:**
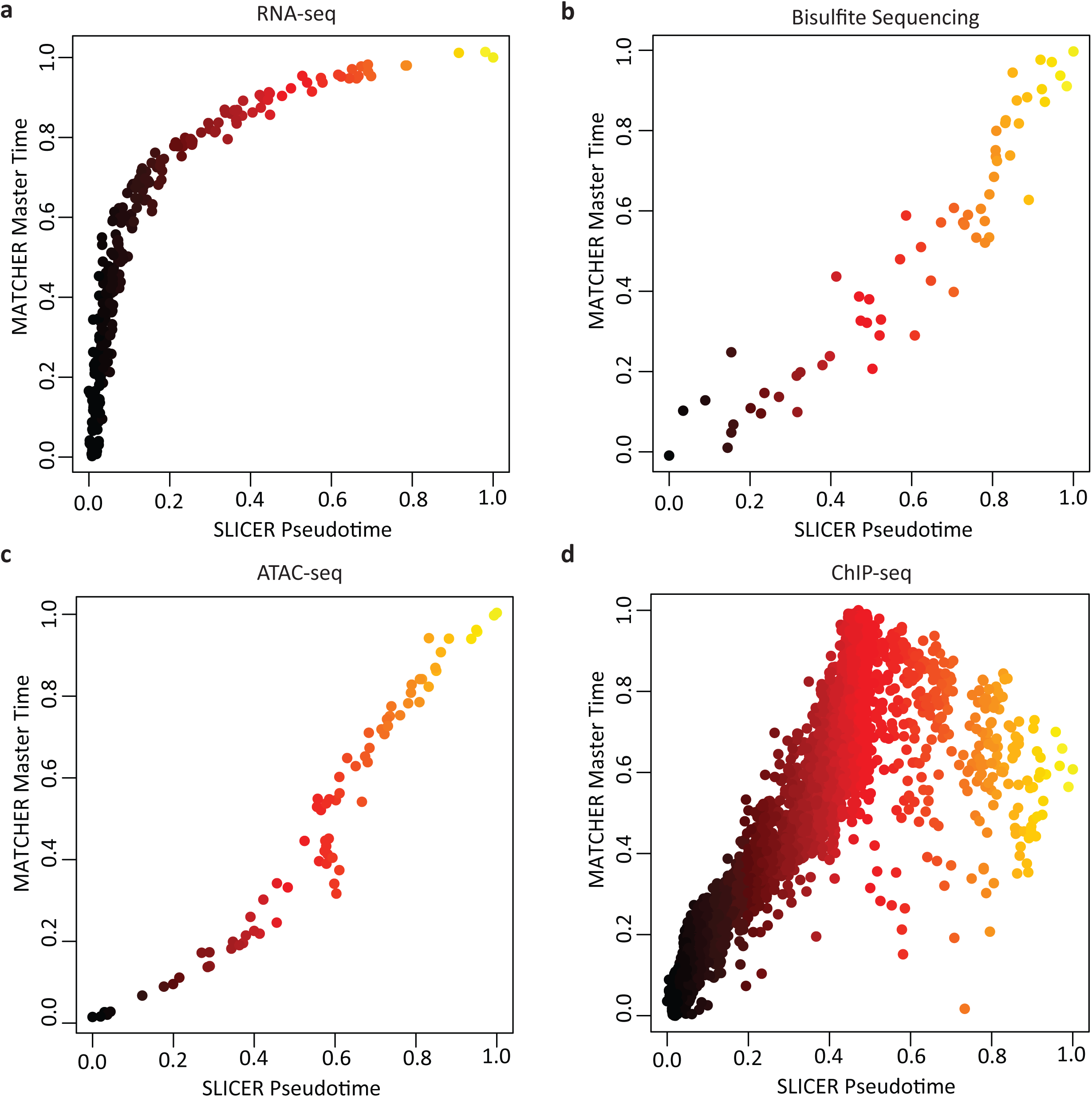
MATCHER master time is strongly correlated with SLICER pseudotime. Scatterplot of SLICER pseudotime versus MATCHER master time for (a) RNA-seq, (b) bisulfite sequencing, (c) ATAC-seq, and (d) H3K4me2 ChIP-seq. The points are colored by SLICER pseudotime.

**Supplementary Figure 2:**
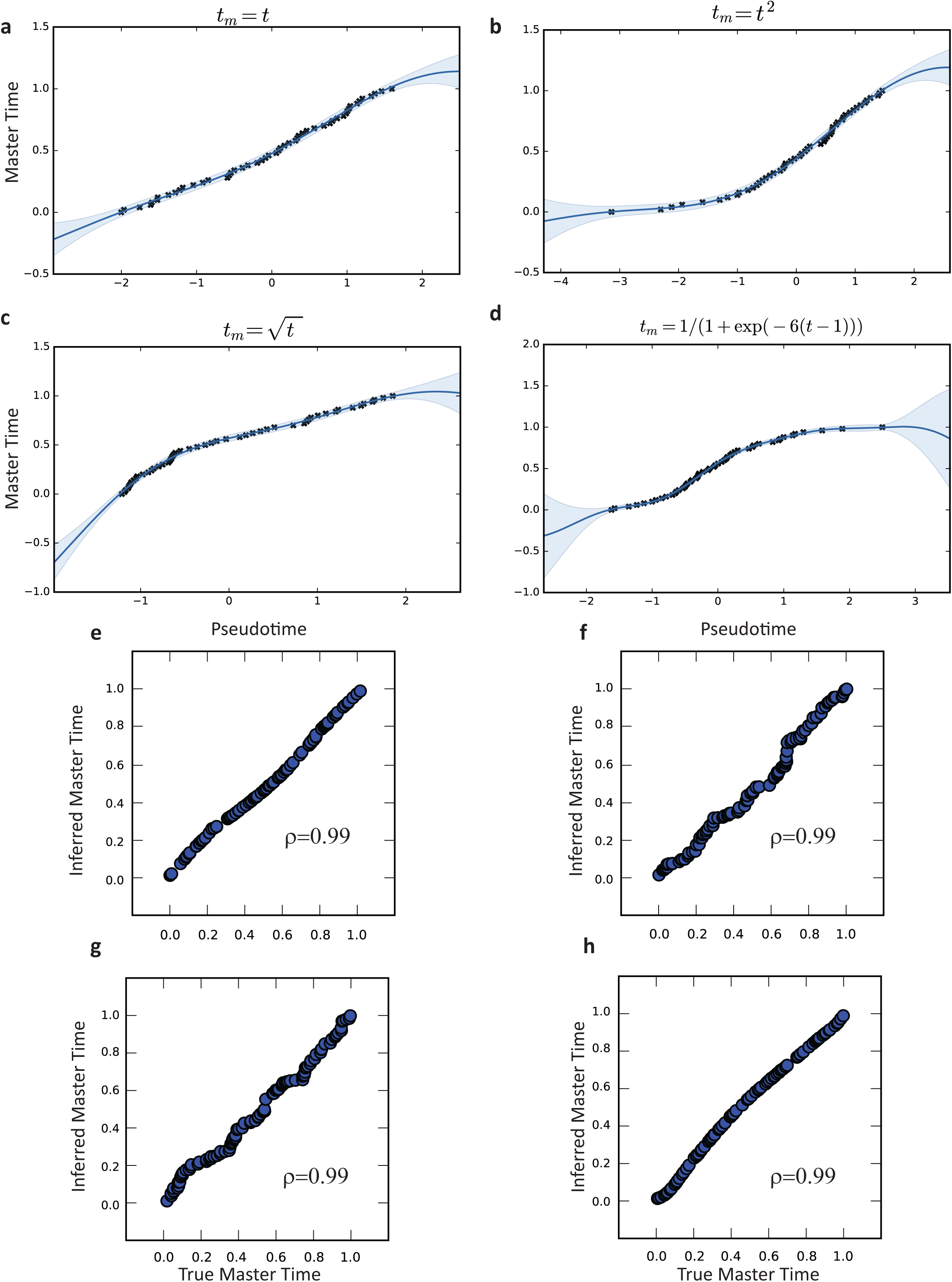
Results from synthetic data generated from different underlying warping functions. Inferred warping functions for (a) linear, (b) square root, (c) quadratic, and (d) logit true underlying warping functions. (e)-(h) Scatterplot of true vs. inferred master time for the corresponding warp functions of panels (a)-(d).

**Supplementary Figure 3:**
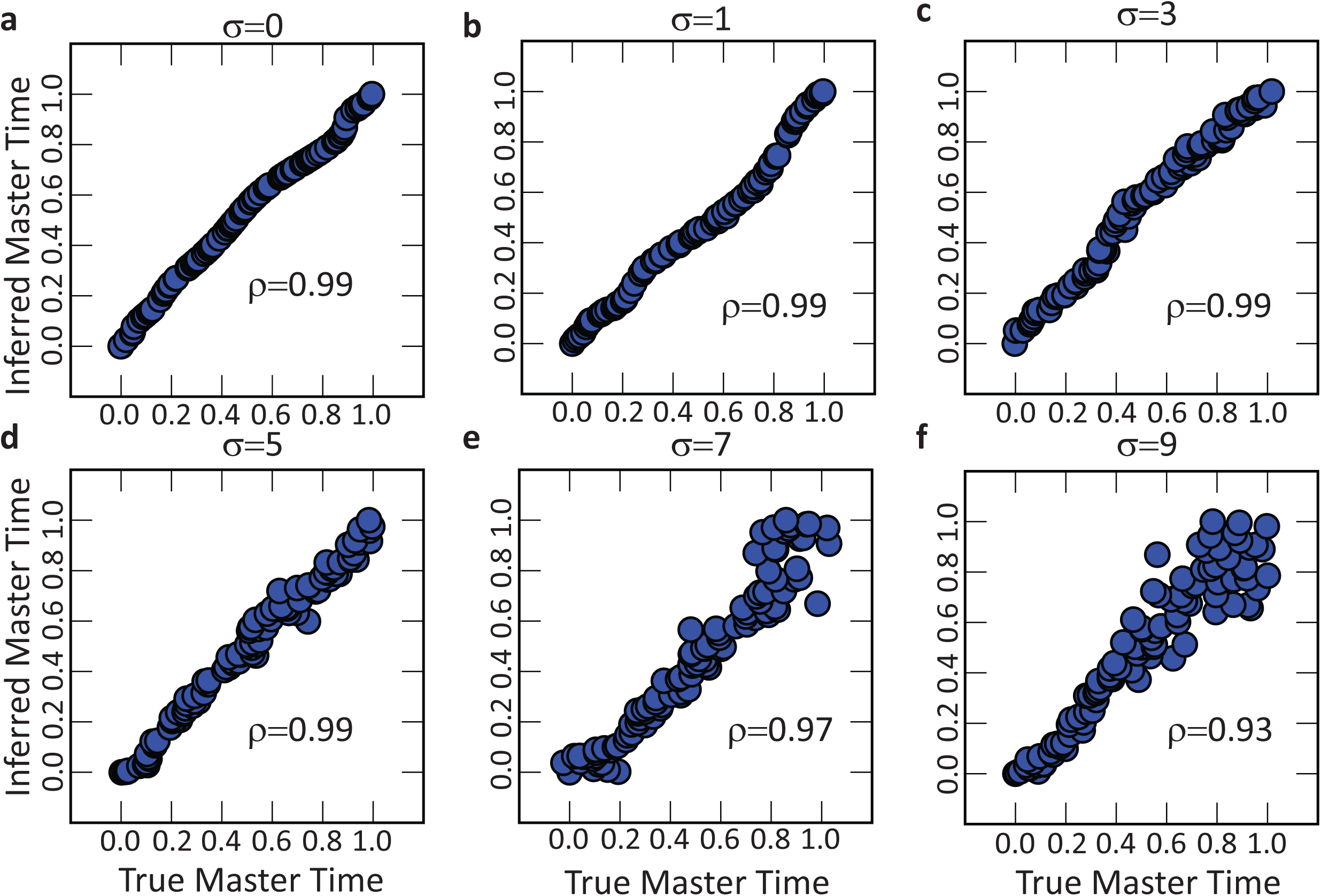
Synthetic data results for increasing noise levels.

**Supplementary Figure 4:**
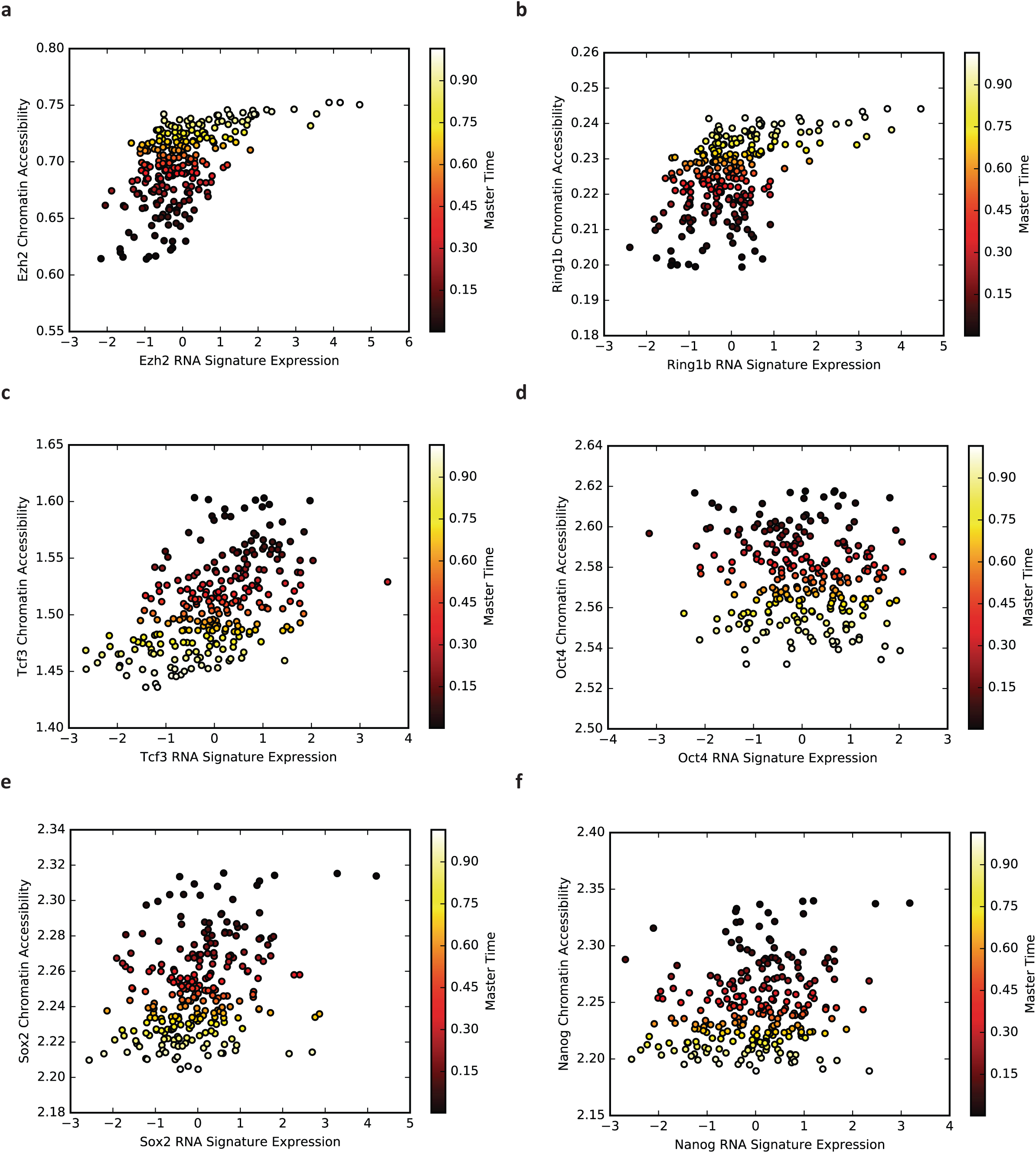
Corresponding values inferred by MATCHER for gene expression and chromatin accessibility signatures. Each point represents inferred correspondence from a single cell. The x-axis shows the value of the gene expression signature in that cell, and the y-axis shows the value of the chromatin accessibility signature. The points are colored by inferred master time. Note that these are the data used to generate the values on the diagonal of the heatmap in Fig. 4c.

**Supplementary Figure 5:**
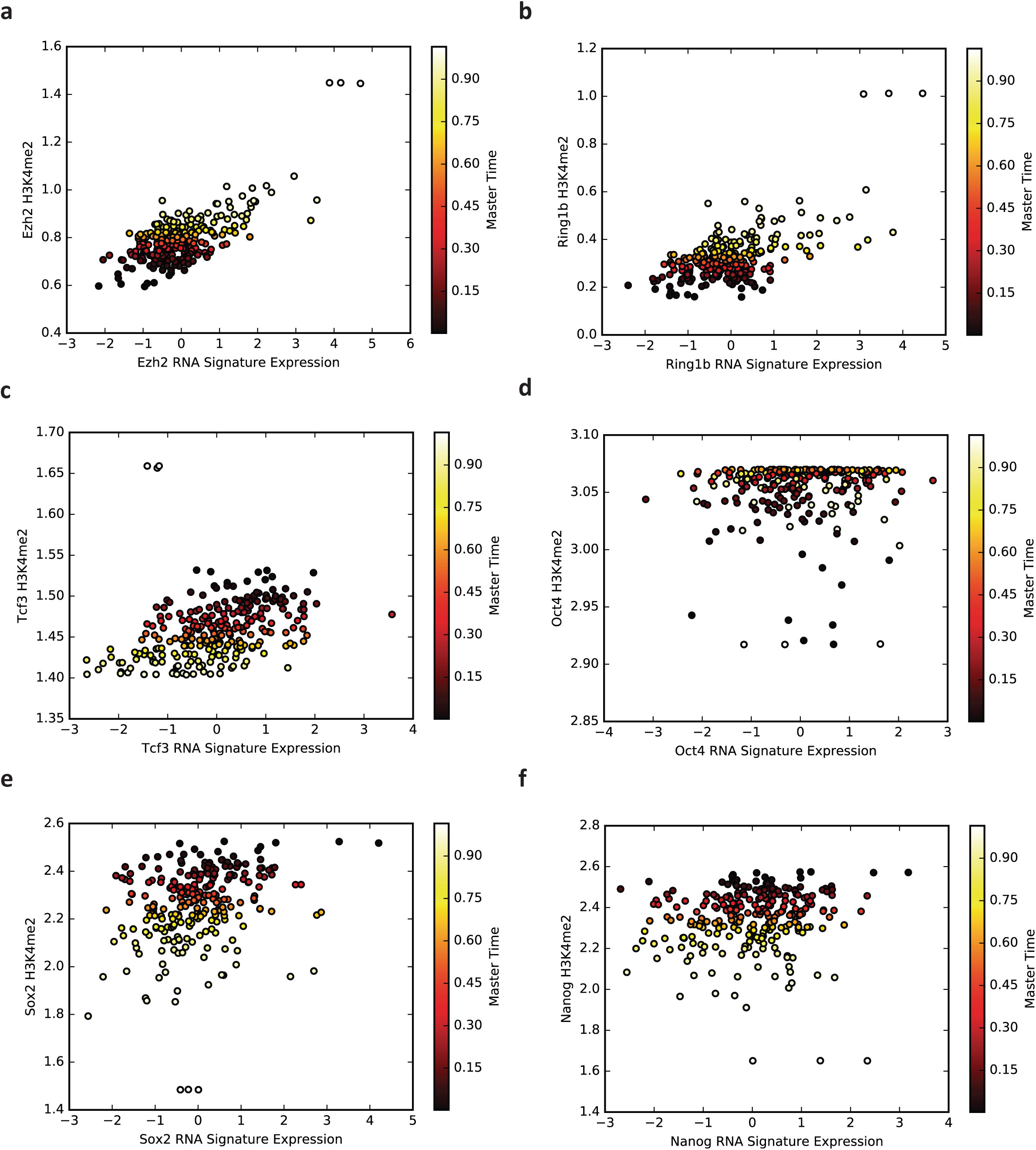
Corresponding values inferred by MATCHER for gene expression and H3K4me2 signatures. Each point represents inferred correspondence from a single cell. The x-axis shows the value of the gene expression signature in that cell, and the y-axis shows the value of the H3K4me2 signature. The points are colored by inferred master time. Note that these are the data used to generate the values on the diagonal of the heatmap in Fig. 4d.

**Supplementary Figure 6:**
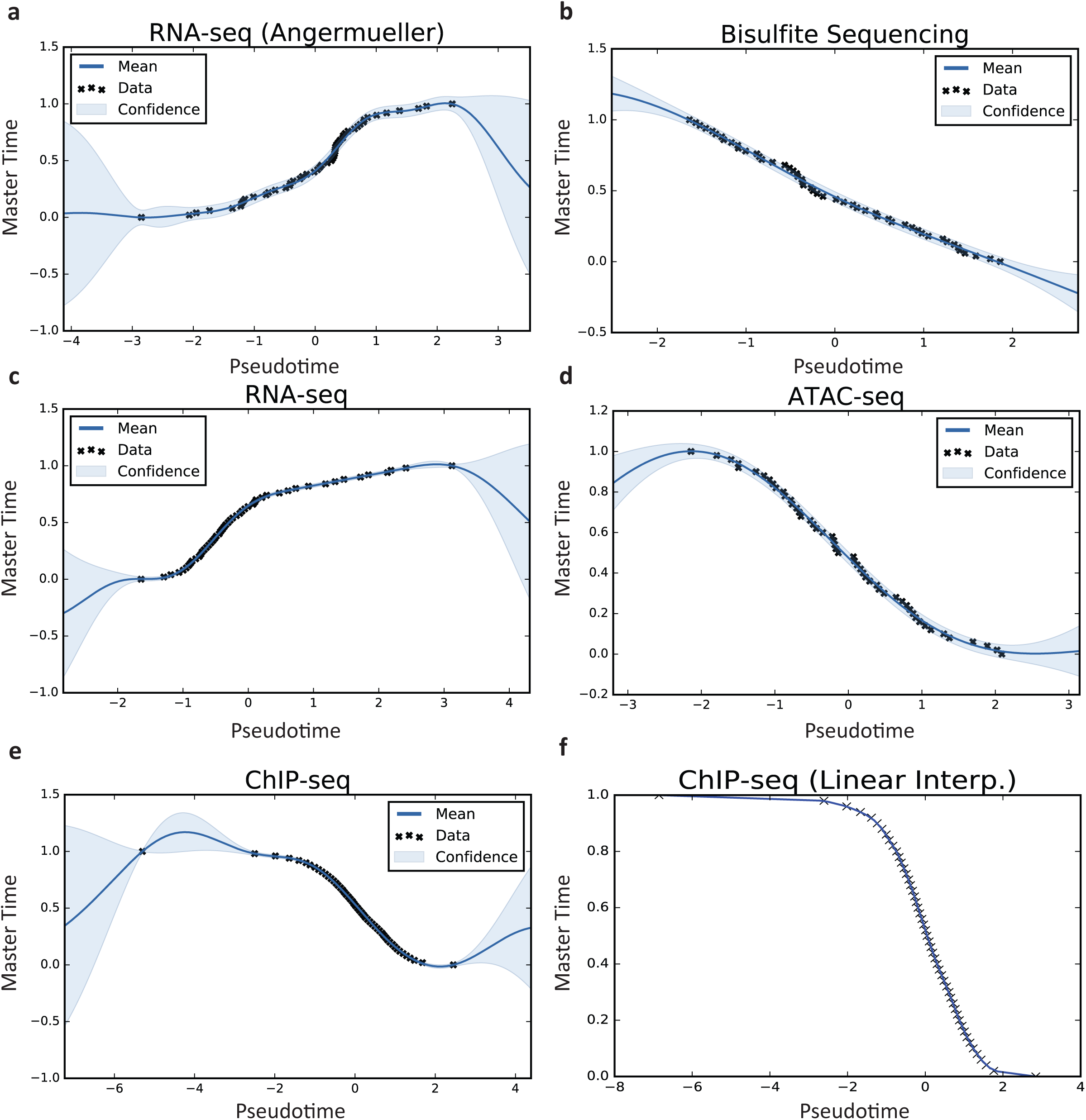
Inferred warping functions for all experimental datasets analyzed in the paper. (a) Kolodziejczyk single cell RNA-seq data, (b) Angermueller scM&T-seq methylation data, (c) ATAC-seq, (d) H3K4me2 ChIP data, (e) Angermueller scM&T-seq gene expression data, and (f) warping function resulting from linear interpolation of H3K4me2 ChIP-seq data.

